# RNF185 control of COL3A1 expression limits prostate cancer migration and metastatic potential

**DOI:** 10.1101/2023.06.29.547118

**Authors:** Benjamin Van Espen, Htoo Zarni Oo, Colin Collins, Ladan Fazli, Alfredo Molinolo, Rabi Murad, Martin Gleave, Ze’ev A. Ronai

**Affiliations:** Cancer Center, Sanford Burnham Prebys Medical Discovery Institute, La Jolla, CA 92037, USA; Vancouver Prostate Centre, Department of Urologic Sciences, University of British Columbia, Vancouver, BC V6H 3Z6, Canada; Department of Pathology, The University of California San Diego, La Jolla CA, 92037

**Keywords:** ubiquitin ligase, RNF185, prostate adenocarcinoma, metastasis, epithelial-mesenchymal transition (EMT), collagen

## Abstract

RNF185 is a RING finger domain-containing ubiquitin ligase implicated in ER-associated degradation. Prostate tumor patient data analysis revealed a negative correlation between RNF185 expression and prostate cancer progression and metastasis. Likewise, several prostate cancer cell lines exhibited greater migration and invasion capabilities in culture upon RNF185 depletion. Subcutaneous inoculation of mouse prostate cancer MPC3 cells stably expressing shRNA against RNF185 into mice resulted in larger tumors and more frequent lung metastases. RNA-sequencing and Ingenuity Pathway Analysis identified wound healing and cellular movement among the most significant pathways upregulated in RNF185-depleted, compared to control prostate cancer cells. Gene Set Enrichment Analyses performed in samples from patients harboring low RNF185 expression and in RNF185-depleted lines confirmed the deregulation of genes implicated in EMT. Among those, COL3A1 was identified as the primary mediator of RNF185’s ability to impact migration phenotypes. Correspondingly, enhanced migration and metastasis of RNF185 KD prostate cancer cells were attenuated upon co-inhibition of COL3A1. Our results identify RNF185 as a gatekeeper of prostate cancer metastasis, partly *via* its control of COL3A1 availability.

## Introduction

Prostate cancer (PCa) is among the most common solid malignancies in developed regions and the second most common in men worldwide^1^. Most PCa diagnosed early in life are not life-threatening; active monitoring is usually the recommended course of action. Nevertheless, a sizeable fraction of primary PCa progresses to advanced disease, as such, PCa is the second leading cause of cancer death in American men^2^. Advanced PCa often harbors an amplification and/or mutation of the androgen receptor (AR), making Androgen Deprivation Therapy (ADT) the primary treatment regime. However, most treated PCa eventually develop resistance to ADT and progress to metastatic disease, known as metastatic Castration-Resistant Prostate Cancer (mCRPC). Once tumors become castration-resistant, the prognosis is bleak, as this form of PCa is considered incurable^3^. Disease trajectory can be somewhat predicted based on histopathological and molecular profiling of tumors. Nonetheless, finding reliable markers of prostate cancer progression and therapeutic targets for mCRPC remains a major unmet clinical need^4^.

Ubiquitination is a post-translational modification in which a ubiquitin moiety is added to lysine residues of target proteins in a process catalyzed by an E1 ubiquitin-activating enzyme, an E2 ubiquitin-conjugating enzyme, and a dedicated E3 ubiquitin ligase. E3 ligases selectively interact with substrates, enabling the formation of ubiquitin chains of different topologies^5^. Those modifications can alter protein localization and complex formation or result in proteasome-mediated degradation of the target protein. Protein ubiquitination enables timely control of key cellular processes, including development, proliferation, cell cycle control, transcription, DNA damage repair, metabolism, and protein-protein interaction. Dysregulation of the ubiquitin system is often implicated in birth defects, neurodegenerative disease, and tumorigenesis^6, 7^. Notably, the impact of E3 ligase dysregulation is highly context-dependent. For example, increased expression of the ubiquitin ligase RNF5 is seen in several human cancers but has different outcomes for breast cancer, melanoma, or acute myeloid leukemia^8–10^.

Growing evidence points to a role for the RNF5-closely related ubiquitin ligase RNF185 in endoplasmic reticulum-associated degradation (ERAD), a process essential for cellular protein homeostasis, which is part of the greater ER stress response. Notably, RNF185 is implicated in cellular functions linked to RNF5^11–17^. The two proteins share >70% sequence identity, and, as seen for RNF5, RNF185 dysregulation reportedly impacts diverse cancer types^18, 19^. However, RNF185 remains poorly characterized and its role in cancer is unclear.

Proteins of the collagen family reportedly play critical roles in diverse cellular processes, including cell adhesion, migration, differentiation, and proliferation^21^. Correspondingly, collagen dysregulation is implicated in several pathologies, including cancer^22–24^. COL3A1 (collagen type III alpha 1) is one of the main collagen types expressed in stromal cells and is mainly produced by fibroblasts, as such, it contributes to regulating the tumor microenvironment (TME)^23, 25^. COL3A1 mRNA expression is upregulated in various cancer subtypes, including breast, lung, head and neck, pancreatic, ovarian, gastric, esophageal cancers, and glioma^26–31^. Moreover, in many of these cases, high COL3A1 expression is associated with increased tumor growth, invasion, metastasis, and sometimes therapy resistance^32–36^.

Here we identify an important function for RNF185 in PCa migration and metastatic behavior and report that COL3A1 mediates these effects. RNF185-depleted prostate cancer cells exhibited enhanced COL3A1 mRNA expression, and in specimens from prostate cancer patients, high COL3A1 expression was correlated with poor prognosis. We also demonstrate that decreasing RNF185 levels increased prostate cancer cell metastatic capacity. Moreover, RNF185 knockdown induced upregulation of COL3A1, which in turn enhanced prostate cancer cell migration and invasion *in vitro* and *in vivo*. We further established that silencing COL3A1 was sufficient to attenuate the increased migration and metastases seen upon RNF185 depletion. Overall, our findings establish RNF185 as functioning in the regulation of COL3A1 levels, an activity that impacts the progression and metastasis of prostate cancer.

## Material & Methods

### Animal studies

All mouse studies were approved by Sanford Burnham Prebys (SBP) IACUC protocol # 20-064. The xenograft model was established using MPC3 cells expressing either control shRNA or shRNA targeting RNF185, COL3A1 or both using the PLKO-1 vector. NOD/SCID/Gamma (NSG) mice were obtained from the SBP Animal Facility. Eight-week-old male mice were injected subcutaneously in the flank with 5 × 10^5^ cells. Mice were housed four per cage and maintained under controlled temperature (22.5°C) and illumination (12 h dark/light cycles) conditions. Mice were sacrificed upon signs of morbidity resulting from tumor growth or if maximal tumor size/burden (2 cm^3^) was reached. Tumor volume was measured with linear calipers and calculated using the following formula: (length in mm × width in mm) × 1/2. After mice were sacrificed, tumors and lungs were snap-frozen and later utilized for histopathological analysis. Fixed and embedded lung tissues were sliced into 5-μm sections to obtain nine serial sections per lung. Metastases in the lungs were evaluated by hematoxylin and eosin staining and quantified by measuring the areas affected by metastases in at total of 9 serial sections for each mouse, using ImageScope v12.3.3 (APERIO, RRID: SCR_014311). The average metastatic area (mm^3^) per section was reported for each group.

### Cell lines and cell culture

Hek293T human kidney epithelial cells (ATCC Cat# CRL-3216, RRID: CVCL_0063) and PCa lines PC3 (ATCC Cat# CRL-1435, RRID: CVCL_0035), C4-2B (ATCC Cat# CRL-3315, RRID:CVCL_4784), TRAMP-C2 (ATCC Cat# CRL-2731, RRID:CVCL_3615) and MyC-CaP (ATCC Cat# CRL-3255, RRID: CVCL_J703)) were obtained from the American Type Culture Collection (ATCC). MPC3 cells were kindly provided by Prof. Zhenbang Chen. PC3 and C4-2B cells were cultured in RPMI growth medium [RPMI supplemented with 10% fetal bovine serum (FBS; Omega Scientific) and 100 units penicillin/mL and 100 units streptomycin/mL]. Hek293T and the mouse PCa line MPC3, TRAMP-C2 and MyC-CaP were cultured in DMEM growth medium (DMEM supplemented with 10% FBS and 100 units penicillin/mL and 100 units streptomycin/mL). Cells were maintained at all times in growth phase. All cell lines were tested negative for mycoplasma contamination with MycoAlert® Mycoplasma Detection Kit (Lonza, LT07-118).

### RNA isolation and quantitative reverse transcription PCR (RT-qPCR)

Total RNA was extracted from cells using a GenElute RNA purification kit (Sigma-Aldrich, RTN70). RNA purity and concentration were assessed using a NanoDrop spectrophotometer (Thermo Fisher). Aliquots of 1 ug of total RNA were reverse transcribed using qScript cDNA synthesis kit (Quantabio, 95047-500) according to the manufacturer’s protocol. qRT-PCR analyses were performed using LightCycler® 480 SYBR Green I Master RT-PCR kits (ROCHE) on a Bio-Rad CFX Connect Real-Time system (Bio-Rad, CFX Connect Real-Time System). Expression levels were normalized to GAPDH. Sequences of oligonucleotide primers were as follows; mouse Rnf185 forward (5′-GCAGACCTATCAATCAACGCC-3’) and reverse (5′-CGAAGGCCCTTTACTTGCCAT-3′); mouse Gapdh forward (5′-AGGTCGGTGTGAACGGATTTG-3’) and reverse (5′-TGTAGACCATGTAGTTGAGGTCA-3′); mouse Col3a1 forward (5′-CTGTAACATGGAAACTGGGGAAA-3’) and reverse (5′-CCATAGCTGAACTGAAAACCACC-3′); mouse Col5a2 forward (5′-TTGGAAACCTTCTCCATGTCAGA-3’) and reverse (5′-TCCCCAGTGGGTGTTATAGGA-3′); human RNF185 forward (5′-GGAGAATAGAGGGGGATTTC-3’) and reverse (5′-AAATGCTGTGGCAAATAT-3′); human GAPDH Forward (5′-GGAGGGAGATCCCTCCAAAAT-3’) and reverse (5′-GGCTGTTGTCATACTTCTCATGG-3′); human COL3A1 forward (5′-GGAGCTGGCTACTTCTCGC-3’) and reverse (5′-GGGAACATCCTCCTTCAACAG-3′); human COL5A2 forward (5′-GACTGTGCCGACCCTGTAAC-3’) and reverse (5′-CCTGGACGACCACGTATGC-3′).

### Western blot analysis

Cells were washed twice with PBS at room temperature and resuspended in lysis buffer [100 mM Tris-HCl pH 7.5, 5% sodium dodecyl sulfate (SDS)]. Lysates were boiled 10 minutes and then sonicated with a Microtip sonicator and subjected to SDS-PAGE. Proteins were transferred to nitrocellulose membranes (Bio-Rad, 1704159). Membranes were blocked in 5% skim milk and incubated with respective primary antibodies followed by HRP-conjugated secondary antibodies (Jackson ImmunoResearch Labs Cat# 111-035-045, RRID: AB_2337938 and Cat# 115-035-062, RRID: AB_2338504). Blots were imaged with a ChemiDoc imaging system (Bio-Rad, ChemiDoc MP imaging system) using ECL Western blotting HRP substrate (Millipore, WBLUR0500).

### Antibodies and chemicals

Antibodies were obtained from the following sources: RNF185 (181999, Abcam, RRID: AB_2922962); β-Tubulin (2128, Cell Signaling Technology, RRID: AB_823664); HRP-conjugated secondary antibodies: goat-anti-mouse-HRP (Jackson ImmunoResearch, Cat# 115-035-062, RRID: AB_2338504) and goat-anti-rabbit-HRP (Jackson ImmunoResearch, Cat# 111-035-045, RRID: AB_2337938). Antibodies were used at a dilution of 1:1000, and blots were washed 3 times with TBST before being incubated with secondary antibodies at a dilution of 1:10000.

### Plasmids, short interfering RNA (siRNA), short hairpin RNA (shRNA) and lentiviral infection

For transient gene depletion, oligonucleotide siRNAs targeting RNF185, COL3A1 and COL5A2, and an siRNA Universal Negative Control were purchased from MISSION® Predesigned siRNA libraries (Sigma-Aldrich). Cells were transfected using JetPRIME transfection reagents (PolyPlus, 101000015). Experiments were performed at 24h or 48h post-transfection. The lentiviral pLKO.1 vector was used to introduce short hairpin RNA (shRNA) targeting RNF185, COL3A1 or the scramble control into MPC3 cells. Sequences for shRNAs were identified through the MISSION® Predesigned shRNA libraries (Sigma-Aldrich) and shRNAs were obtained from the La Jolla Institute for Immunology RNAi Center (La Jolla, CA, USA). pLKO.shRNF185, pLKO.shCOL3A1 and pLKO.scramble infectious lentiviral stocks were generated in Hek293T cells. MPC3 cells were infected 8 h with lentivirus in the presence of 8 μg/mL polybrene (Millipore). After 48 h, cells stably expressing scramble or targeted shRNA(s) were selected in 2 μg/mL puromycin. Five days later, puromycin concentration was halved. Puromycin treatment was stopped 48 h before beginning experimental procedures.

### Cell proliferation assay

Cells were plated in 96-well clear-bottomed plates (Corning, 3610) at a density of 2,000 cells/100 μL of medium per well. Proliferation was measured every 24h by treating cells with 30 μL of the CellTiter Glo kit reagent (Promega) and measuring luminescence with a CLARIOstar Plus microplate reader (BMG Labtech).

### Cell migration and invasion assay (Boyden chamber)

Transwell plates were obtained from Corning (#3342). For migration assays, 2 × 10^4^ cells of the MPC3, MyC-CaP and TRAMP-C2 lines and 6 × 10^4^ cells of PC3 and C4-2B lines were seeded into the upper chamber in 100 μL serum-free medium. The lower chamber was filled with 600 μL of medium supplemented with 10% FBS as a chemo-attractant. After incubations of 24h or 48h (C4-2B), cells were fixed in 4% PFA and stained with 0.5% crystal violet. Cells on the upper surface of the filter were removed with a cotton swab. For quantification, crystal violet stain was eluted by incubating inserts in 30% acetic acid, and absorbance was measured at 492 nm using a CLARIOstar Plus microplate reader (BMG Labtech). For invasion assays, Transwells were first coated with Matrigel overnight, and then procedures described for the migration assay were followed.

### Spheroid formation assay

Cells were plated in Nunc™ MicroWell™ MiniTrays (ThermoFisher # 438733) at a density of 200 cells/20 μL of medium per well. The tray was then flipped upside-down to allow cells to grow in hanging droplets and form 3D spheroids in which cells are in direct contact with each other and with extracellular matrix components. The trays were placed in 150 mm tissue culture dish with 5 mL of PBS and a piece of Whatman filter paper to act as hydration chamber. Spheroids size was measured after 3, 5 and 9 days of incubation. For quantification, pictures of each spheroid were processed with ImageJ (RRID:SCR_003070), where surface areas occupied by spheroids were measured.

### RNA-seq library construction

PolyA RNA was isolated using NEBNext® Poly(A) mRNA Magnetic Isolation Module, and bar-coded libraries were constructed using the NEBNext® Ultra™ Directional RNA Library Prep Kit for Illumina® (NEB, Ipswich MA). Libraries were pooled and sequenced single-end (1X75) on an Illumina NextSeq 500 using the High output V2 kit (Illumina Inc., San Diego CA).

### RNA-seq processing and analysis

Raw sequencing reads were pre-processed to remove llumina Truseq adapters and polyA/polyT sequences using Cutadapt version 2.3^36^. Trimmed reads were aligned against mouse genome version mm10 and Ensembl gene annotations version 84 using STAR version 2.7.0d_0221 (RRID:SCR_004463)^38^, and alignment parameters from ENCODE long RNA-seq pipeline (https://github.com/ENCODE-DCC/long-rna-seq-pipeline). Gene level estimated counts and transcripts per million (TPM) were obtained using RSEM version 1.3.1^39^. FastQC version 0.11.5 (https://www.bioinformatics.babraham.ac.uk/projects/fastqc/; RRID: SCR_014583) and MultiQC version 1.8 were used to assess the quality of trimmed reads and alignment to genome/transcriptome^40^. Low expressed genes were removed from downstream analysis by selecting the genes with RSEM estimated counts equal or greater than 5 times the total number of samples. Batch effect correction was performed using *Combat_seq* function in sva version 3.44.0. The sequencing libraries were built in three batches: batch1 contained replicates 1 and 2 of each condition, batch 2 contained replicates 3 and 4, and batch 3 contained replicates 5 and 6. Differential expression comparisons were performed using batch-corrected counts and Wald test implemented in DESeq2 version 1.22.2 (RRID: SCR_000154)^41^. Genes with Benjamini-Hochberg corrected p-value < 0.05 and fold change ≥ 2.0, or ≤ -2.0 were identified as differentially expressed. Counts per million (CPM) and reads per kilobase per million (RPKM) values were computed from batch-corrected counts using edgeR version 3.38.4 (RRID: SCR_012802)^42^. Gene lengths (sum of exon lengths) for RPKM calculation were computed from Ensembl gene annotations version 84 GTF file using GenomicsFeatures Bioconductor package version 1.48.4 (RRID: SCR_006442)^43^. Gene set enrichment analysis (GSEA) for TCGA-PRAD comparison of RNF185 low versus high expressors was performed using preranked option in GSEA version 4.2.3 (RRID: SCR_003199)^44^. GSEA for RNA-seq comparisons were performed using GSEA version 4.3.2 and RPKM values with parameter “Permutation type = gene_set”. Pathway analysis was performed using Ingenuity Pathway Analysis (Qiagen, Redwood City, USA; RRID: SCR_008653).

### Analysis of clinical databases

Clinical samples from the Taylor et al. prostate adenocarcinoma cohort and The Cancer Genome Atlas (TCGA) Pan-Cancer prostate adenocarcinoma (PRAD) dataset were used^44, 45^. Gene expression and clinical annotation were downloaded from cBioPortal Cancer Genomic Portal (http://cbioportal.org)^46^. The bottom and top quartiles were used to identify mRNA expression higher or lower than normal prostate samples. The Kaplan–Meier method was used for disease-free survival analysis. For comparison of RNF185 and COL3A1 mRNA expression in samples stratified by Gleason score and by sample type (primary vs. metastasis), the log2 (fold-change) value was used.

### Statistical analysis

At least 3 samples were used in each experimental group. All experiments were performed at least three times to establish statistical power and reproducibility. Differences between two groups were assessed using two-tailed unpaired t-tests or Mann-Whitney u test. One-way ANOVA with Tukey’s multiple comparison test was used to evaluate experiments involving one factor but multiple groups. Two-way ANOVA with Tukey’s multiple comparison test was used to evaluate experiments involving two factors and multiple groups. Fisher’s exact test was used to analyze 2x2 contingency table of small sample size. Disease-free survival was analyzed by the Kaplan–Meier method and evaluated with a log-rank (Mantel-Cox) test. All analyses were performed using GraphPad Prism software (RRID: SCR_002798). P < 0.05 was considered significant.

### Data Availability

Data analyzed in this study were obtained from cBioPortal Cancer Genomic Portal (http://cbioportal.org)^46^. Raw and processed RNAseq data have been deposited at Gene Expression Ominibus GEO under accession GSE228999. Raw data generated in this study were generated using immunohistochemistry, Western blots and qPCR analyses, and all are available upon request from the corresponding author.

## Results

### Low RNF185 expression correlates with metastatic load and unfavorable PCa patient outcomes

Analysis of RNF185 mRNA levels in independent cohorts of PCa patients revealed that low RNF185 mRNA expression was associated with poorer disease-free survival in the prostate adenocarcinoma (PRAD) dataset from The Cancer Genome Atlas (TCGA)^45^. Consistently, analysis of an independent patient cohort consisting of ∼218 human prostate tumors (181 primary tumors and 37 metastases) revealed that low RNF185 expression coincided with poorer prognosis^44^. Median disease-free survival was 53.82 months in RNF185-low samples and undefined (survival >50% at the longest time-point) in all others. In the RNF185-low expressing group, 13 of 24 (54.2%) progressed, whereas 19 of 88 (21.6%) progressed in the RNF185-high expressing group (Fig. 1A). Notably, 52.94% of RNF185-low expressing samples were metastatic compared with 5.2% in the RNF185-high expressing group (Fig. 1B). Likewise, metastatic samples showed significantly lower RNF185 mRNA expression than non-metastatic samples (Fig. 1C). To confirm that lower RNF185 mRNA levels are associated with poorer prognosis, we examined RNF185 expression in samples stratified based on Gleason grade, a histopathological measure of prostate tumor stage and aggressiveness. RNF185 expression in high Gleason grade tumors was significantly lower relative to intermediate and low Gleason grade tumors. (Fig. 1D). In TCGA’s PRAD dataset, samples with low RNF185 mRNA levels also correlated with poor disease-free survival, with a median of 45.24 disease-free months compared to a >50% disease-free survival rate associated with higher RNF185 expression. In the RNF185-low expressing group, 12 of 32 (37.5%) progressed compared with 17 of 299 (5.7%) in the RNF185-high expressing group (Fig. 1E). Lower RNF185 mRNA levels were also associated with higher Gleason grade, shorter progression-free survival, increased new neoplastic events post initial therapy, greater lymph node involvement and higher tumor stage in that dataset (Fig. 1F, Supp. Fig. 1A-D).

**Fig. 1.**
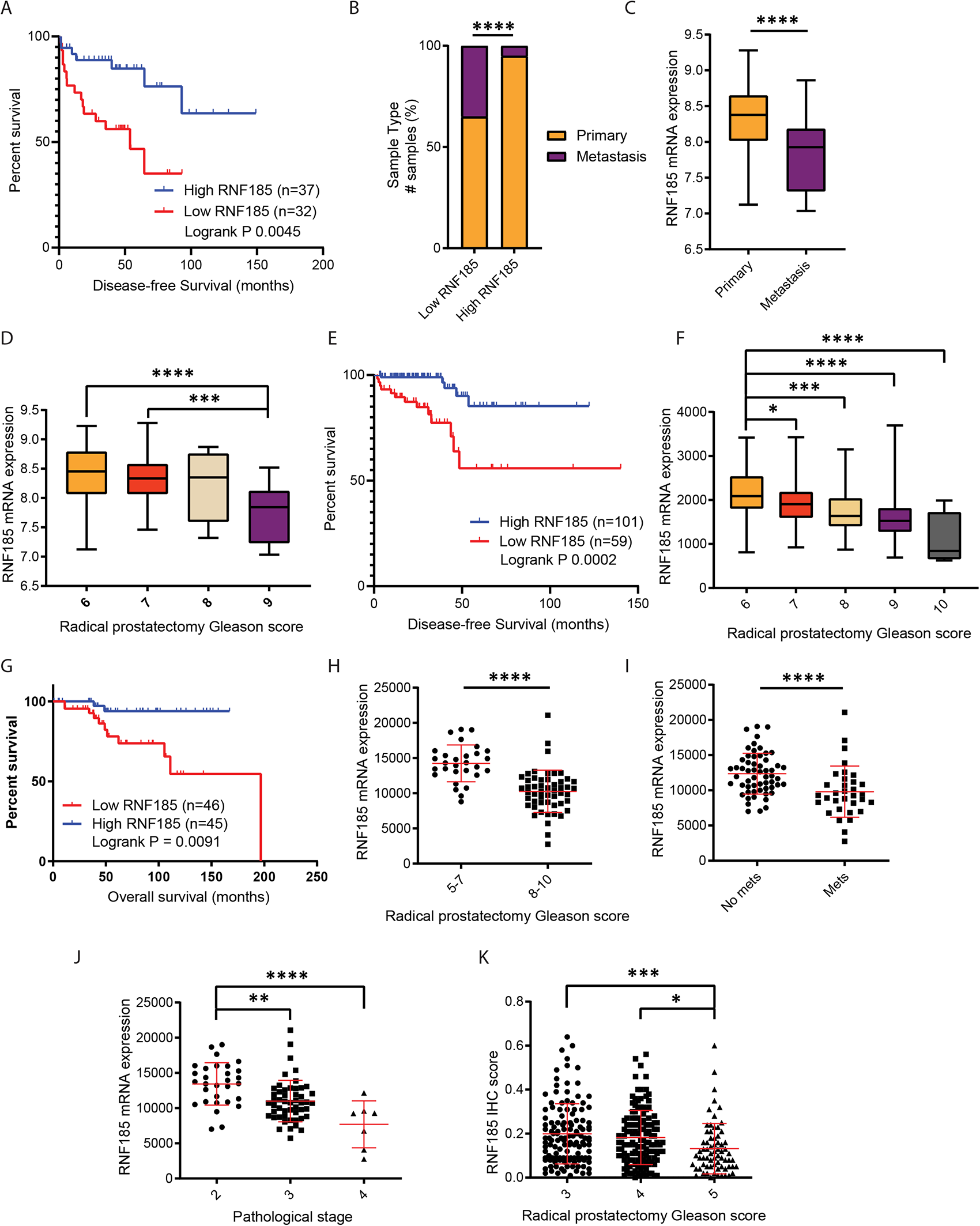
RNF185 expression correlates with metastatic load and PCa patient outcome. (A) Disease-free survival in TCGA-PRAD samples stratified by RNF185 mRNA expression (“Low RNF185” = bottom quartile; “High RNF185” = top quartile). (B) Proportion of sample types in Low RNF185 vs. High RNF185. (C) RNA-seq analysis of RNF185 mRNA in primary tumors (n=131) and metastasis (n=19). (D) RNA-seq analysis of RNF185 mRNA in samples sorted by Gleason score (6: n=41; 7: n=76; 8: n=11; 9: n=11). (E) Disease-free survival in samples stratified by RNF185 mRNA expression using the same cutoffs as in (A). (F) RNA-seq analysis of RNF185 in samples sorted by Gleason score (6: n=45; 7: n=247; 8: n=64; 9: n=138; 10: n=4). (G) Overall survival in PCa samples stratified by RNF185 mRNA expression using the median as cutoffs. (H) RNA-seq analysis of RNF185 in samples sorted by Gleason score (5-7: n=29; 8-10: n=52). (I) RNA-seq analysis of RNF185 in patients with (n=33) and without (n=58) metastases. (J) RNA-seq analysis of RNF185 in samples sorted by pathological stages (2: n=30; 3: n=51; 4: n=7). (K) RNF185 IHC staining score in PCa TMA stratified by Gleason groups (3: n=127; 4: n=133; 5: n=69). (A-D) generated using datasets from Taylor et al., PMID: 20579941; (E-F) generated using TCGA’s Pan-Cancer Atlas and Firehose legacy PRAD datasets. (G-K) generated from VPC unpublished cohorts. Box plots show the median and whiskers (min to max), scatter plots show mean with SD. *P < 0.05, **P < 0.005, ***P < 0.001, ****P < 0.0001 by two-tailed t-test, Mann-Whitney u test, one-way ANOVA or Fisher’s exact test (B). Survival graphs used the Kaplan–Meier with a log-rank (Mantel-Cox).

Further assessment that was carried out using an independent patient cohort (VPC) revealed a negative correlation between RNF185 expression and overall survival (Fig. 1G). Lower RNF185 expression was confirmed in patients with higher Gleason grades (Fig. 1H), with metastases (Fig. 1I), and with more advanced pathological stages (Fig. 1J). Overall, patients considered at high risk exhibited lower RNF185 mRNA expression (Supp. Fig. 1E).

Given that RNF185 is a ubiquitin ligase, its abundance, subcellular localization, and/or activity could be altered by self-degradation and post-translational modification. Therefore, RNF185 transcript levels may not reflect its activity. Evaluation of RNF185 protein levels by immunohistochemistry (IHC) staining of a tissue microarray (TMA) derived from PCa patients revealed a significant decrease of RNF185 staining intensity in samples assigned with the highest Gleason groups: the RNF185 IHC score was significantly lower in samples from Gleason group 5 (high risk) relative to samples from Gleason groups 3 and 4 (intermediate risk) (Fig.1K). Weaker RNF185 staining intensity also correlated with higher percentage of recurrence (Supp. Fig. 1F). These observations, using 3 independent patient cohorts, establish that RNF185 expression (at both transcript and protein levels) inversely correlates with advanced PCa stage (Gleason grade), propensity to metastasize and decreased overall survival.

### RNF185 KD enhances PCa cell migration, tumor development and metastatic load

The relationship between RNF185 mRNA levels and unfavorable patient outcomes prompted us to test the effect of altering RNF185 expression in PCa-derived cell lines, both *in vitro* and *in vivo*. To this end, we infected the mouse PCa line MPC3 (*Pten^-/-^::p53^-/-^*)^47, 48^ with lentivirus expressing control shRNA or shRNF185. MPC3 cells stably expressing shRNF185, confirmed to express low levels of RNF185 (RNF185 KD), were selected (Fig. 2A), and monitored for potential changes in migration and invasion. RNF185 KD MPC3 cells exhibited significantly greater migration and invasion capacity compared with control shRNA cells (Fig. 2B-C). In contrast, RNF185 KD did not alter proliferation of cells, grown under two-dimensional conditions (Fig. 2D). Interestingly, the effect of RNF185 KD on migration seen in MPC3 cells were not observed in mouse prostate cancer cell lines TRAMP-C2 and MyC-CaP, which harbor different mutations^49, 50^ (Supp. Fig. 2A-F), implying that select genetic drivers of PCa could impact the ability of RNF185 to alter PCa phenotypes.

**Fig. 2.**
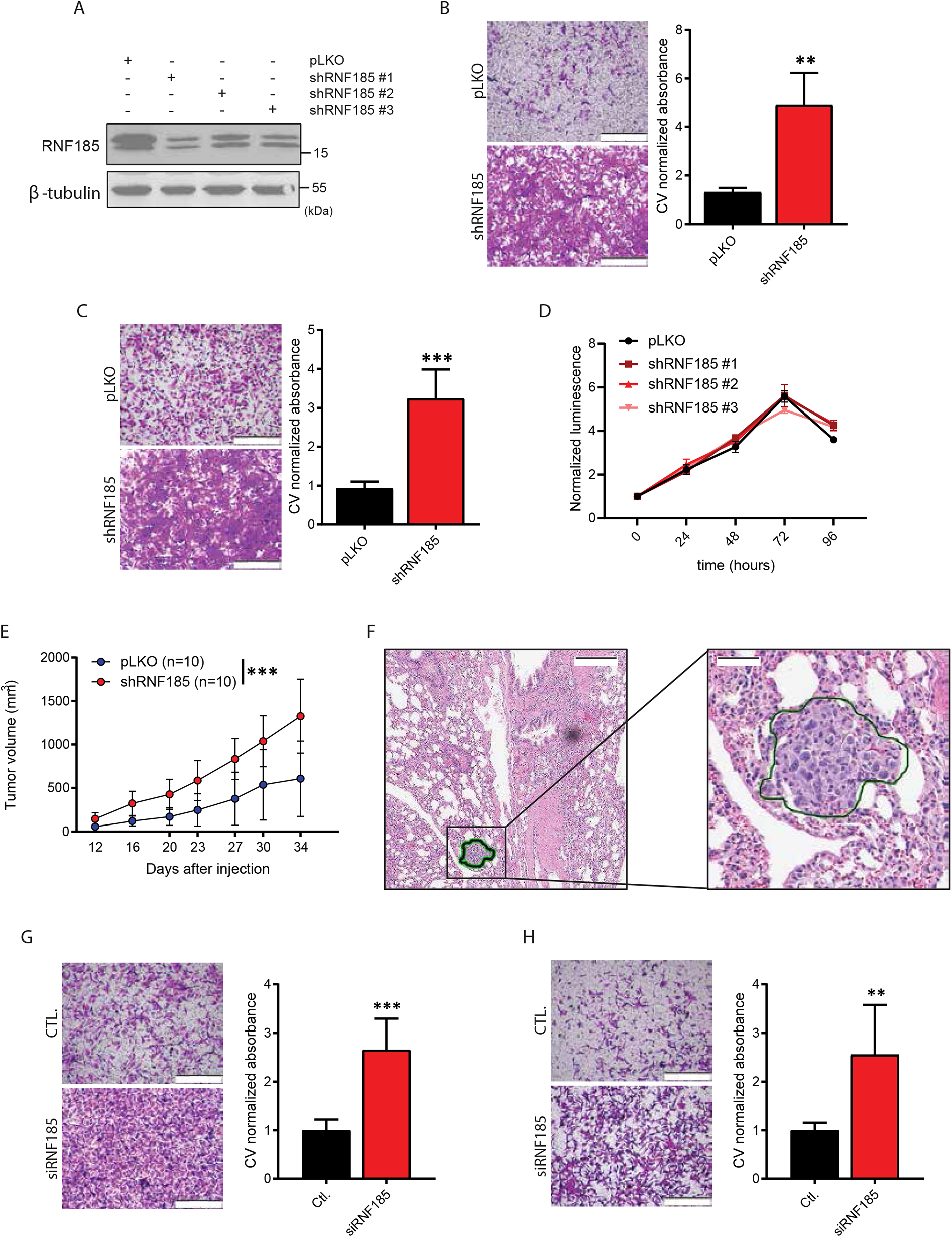
RNF185 KD enhances PCa migration, tumor development and metastatic load. (A) Western blot analysis of RNF185 protein levels in lysates of MPC3 cells stably expressing the indicated vectors. (B) Representative pictures of migrated MPC3 cells expressing the indicated vectors and quantification of migrated cells based on Crystal Violet (CV) absorbance. (C) Representative images of MPC3 cells expressing the indicated vectors that migrate through Matrigel and quantification of those migrated cells based on Crystal Violet absorbance. (D) Proliferation assay of the same cells shown in (A). (E) Growth of MPC3-pLKO and MPC3-shRNF185 cells after subcutaneous injection of 5x10^5^ cells into the flank of NSG mice (n=10/group). (F) Representative H&E staining of lung metastasis from mice inoculated with MPC3-shRNF185 cells (34 days post-injection). (G) Representative images of migrated PC3 cells transfected with siRNF185 or Universal negative control and quantification of migrated cells based on Crystal Violet (CV) staining absorbance. (H) Representative images of migrated C4-2B cells transfected with siRNF185 or Universal negative control and quantification of migrated cells based on Crystal Violet (CV) absorbance. Box plots show the median and whiskers (min to max). Growth curve shows the mean and whiskers (min to max). Box plots show median and whiskers. **P < 0.005, ***P < 0.001, ****P < 0.0001 by two-tailed t-test or two-way ANOVA.

To monitor possible changes in MPC3 cell metastatic propensity *in vivo*, we inoculated NSG mice subcutaneously with RNF185 KD or control cells. Larger tumors were observed in mice inoculated with RNF185 KD as compared to WT MPC3 cells (Fig. 2E, Supp. Fig. 2H). Notably, 7 of 20 mice inoculated with RNF185 KD cells developed lung metastases (a total of 13 micro-metastases were detected) compared with 1 of 20 in the control MPC3 tumor cohort (only 1 micro-metastasis detected). These observations are consistent with enhanced migration phenotypes seen in cultured MPC3 cells (Fig. 2F). Enhanced migratory capability upon RNF185 knock-down was validated in cultures of human prostate cancer lines PC3 and C4-2B (Fig. 2G-H). Both lines also formed larger spheroids upon RNF185 inhibition (Supp. Fig. 2I-J). These analyses demonstrate that inhibition of RNF185 expression is sufficient to increase the metastatic potential of select prostate cancer cells.

### Transcriptional analysis suggests activation of the Epithelial-to-Mesenchymal Transition and collagen upregulation in RNF185 KD cells

To identify possible mechanisms underlying RNF185 effect on cell migration, we performed RNA-seq analysis of RNF185 KD and control MPC3 lines. Gene expression analysis was performed using six replicate samples of MPC3 cells expressing one of two different shRNAs (shRNF185 #1 and #3) compared with control shRNA-infected cells. Analysis of differentially-expressed genes (DEGs) revealed clustering that distinguished the control from the two RNF185 KD lines (Fig. 3A). Even though shRNF185#1 and shRNF185#3 clustered separately in the Principal Component Analysis, Ingenuity Pathway Analysis (IPA) yielded the same results in both groups when compared to controls. In both cases IPA identified gene clusters representing cellular movement/migration as upregulated, coinciding with phenotypic changes seen in RNF185 KD MPC3 cells (Fig. 3B). Furthermore, there was a significant overlap in up-or down-regulated genes between RNF185 KD lines #1 and #3 relative to controls (Supp. Fig. 3A). To identify potential drivers of migration phenotypes, we compared MPC3 cell RNA-seq data with clinically available transcriptomics data. Gene set enrichment analysis (GSEA) was first performed using differentially expressed gene lists from TCGA’s PRAD dataset in which patients were clustered based on RNF185 mRNA expression (bottom 25% vs. top 25%). Next, we carried out GSEA analysis based on RNA-seq of RNF185-depleted MPC3 cells. Analysis of the Molecular Signatures Database (MSigDB) identified a hallmark gene set enriched for factors associated with the Epithelial-to-Mesenchymal Transition (EMT) (Fig. 3C, Supp. Table 1-3)^51^. A list of select genes driving the enrichment score and common to all datasets was generated and used for independent validation (Fig. 3D-E, Supp. Fig. 3B). This list included a number of shared genes encoding fibrillar collagens that were identified as potential drivers. Among these, upregulation of COL3A1 and COL5A2, upon RNF185 KD, was validated in both mouse and human lines. These collagen genes were therefore selected for further evaluation (Fig. 3E).

**Fig. 3.**
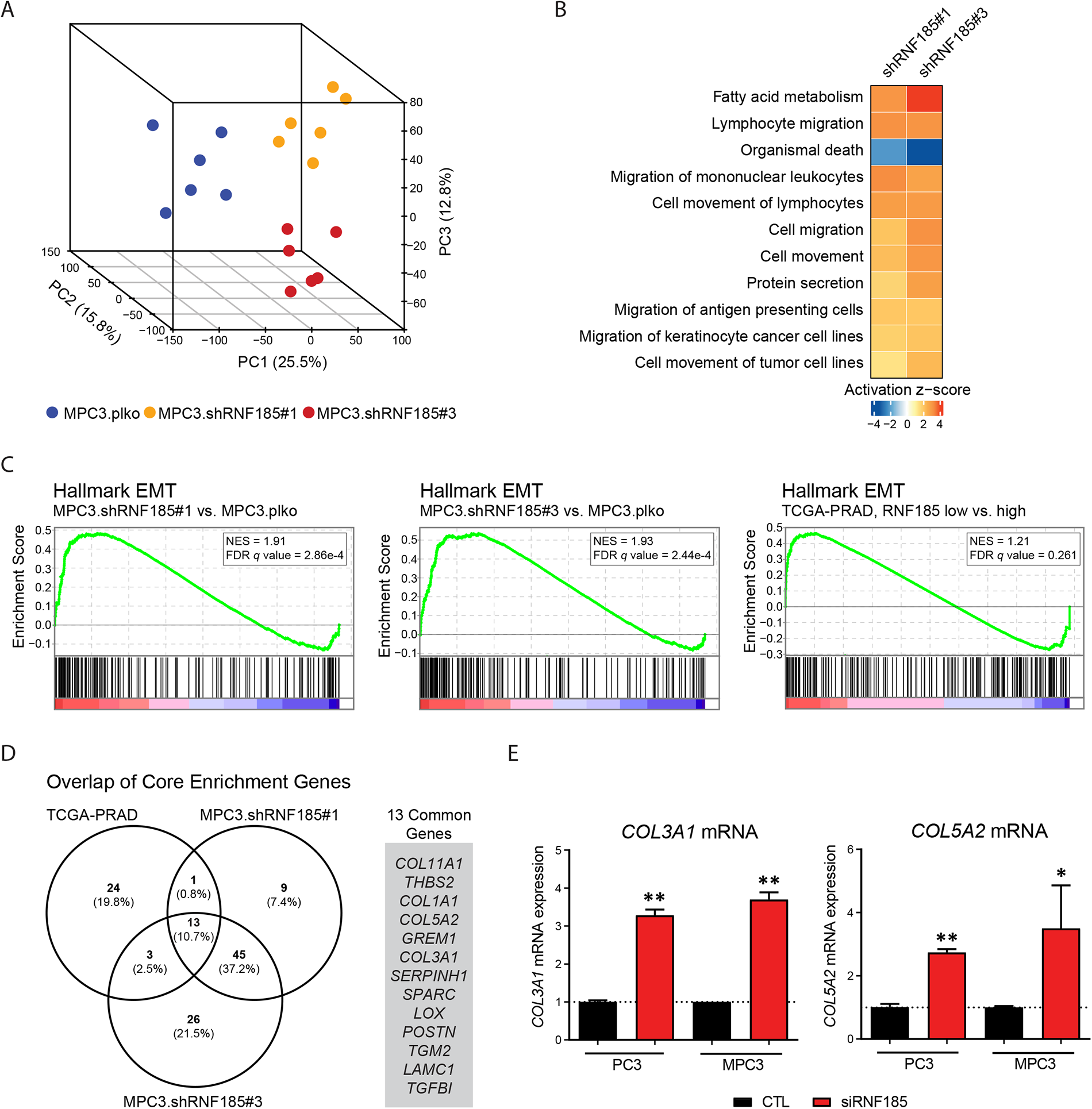
Transcriptional analysis of RNF185 KD MPC3 cells and comparison analysis with clinical transcriptomics reveals EMT pathway activation and collagen upregulation. (A) Principal Component Analysis (PCA) plot of MPC3-pLKO and MPC3-shRNF185 #1 and #3 biological replicates. Heatmap comparing significantly altered diseases and functions in MPC3-shRNF185 #1 and #3 compared to MPC3-pLKO. (C) GSEA enrichment plots showing enrichment of the EMT hallmark gene set in MPC3-shRNF185 #1 and #3 relative to MPC3-pLKO, and in TCGA’s PRAD patients with low RNF185 mRNA expression (bottom 25%) relative to those with high expression (top 25%). (D) Venn Diagrams of enriched genes from HALLMARK_EMT in MPC3-shRNF185 #1 and #3 cells and in low RNF185 expressing patients from TCGA dataset. (E) qRT-PCR analysis of COL5A2 and COL3A1 in MPC3-pLKO and MPC3-shRNF185 cells, and in the human prostate cancer line PC3 transfected with RNF185 siRNA. Box plots show median and whiskers.

### COL3A1 drives enhanced migration in vitro and in vivo and correlates with unfavorable patient prognosis

To directly assess whether changes seen upon RNF185 KD are mediated by upregulation of COL3A1 or COL5A2, we performed rescue experiments in PC3 cells. Each of these genes were inhibited, using the corresponding siRNA, individually or together with RNF185, followed by monitoring possible changes in cell migration. Inhibition of COL3A1 had no effect on migration, while inhibition of COL5A2 increased migration. Notably, however, inhibition of COL3A1 in RNF185 KD cells was sufficient to attenuate the increased migration seen upon RNF185 KD. Conversely, inhibition of COL5A2 did not affect the migration phenotypes seen following RNF185 KD (Fig. 4A-B). As expected, inhibition of both COL3A1 and COL5A2 in RNF185 KD cells also rescued migration phenotypes, which was seen by KD of COL3A1 alone (Fig. 4A-B). Interestingly, COL5A2 depletion induced COL3A1 upregulation and vice versa (Supp. Fig. 4A-C), suggesting a possible cross talk that links their expression. Consistent with these observations, analysis of patient samples (Taylor et al. dataset) revealed that high COL3A1 expression coincided with poorer prognosis. In the COL3A1-high group, 17 of 33 (51.5%) progressed, whereas only 3 of 36 (8.3%) progressed in the COL3A1-low group (Fig. 4C). COL3A1 expression in high Gleason grade tumors was significantly greater than in intermediate and lower Gleason grade tumors (Supp. Fig. 4D). Along these lines, mRNA expression of RNF185 and COL3A1 were found to be negatively correlated in the TCGA PRAD dataset (Fig. 4D). Analyses of independent PCa datasets (from VPC) revealed that COL3A1 expression was primarily seen in patients with higher Gleason grades (Fig. 4E), with a more advanced pathological stages (Fig. 4F), and with more metastases (Fig. 4G). Overall, patients considered at high risk exhibited higher COL3A1 mRNA expression (Fig. 4H). A positive correlation between COL3A1 expression and PSA recurrence was also observed in this patient cohort (Fig. 4I).

**Fig. 4.**
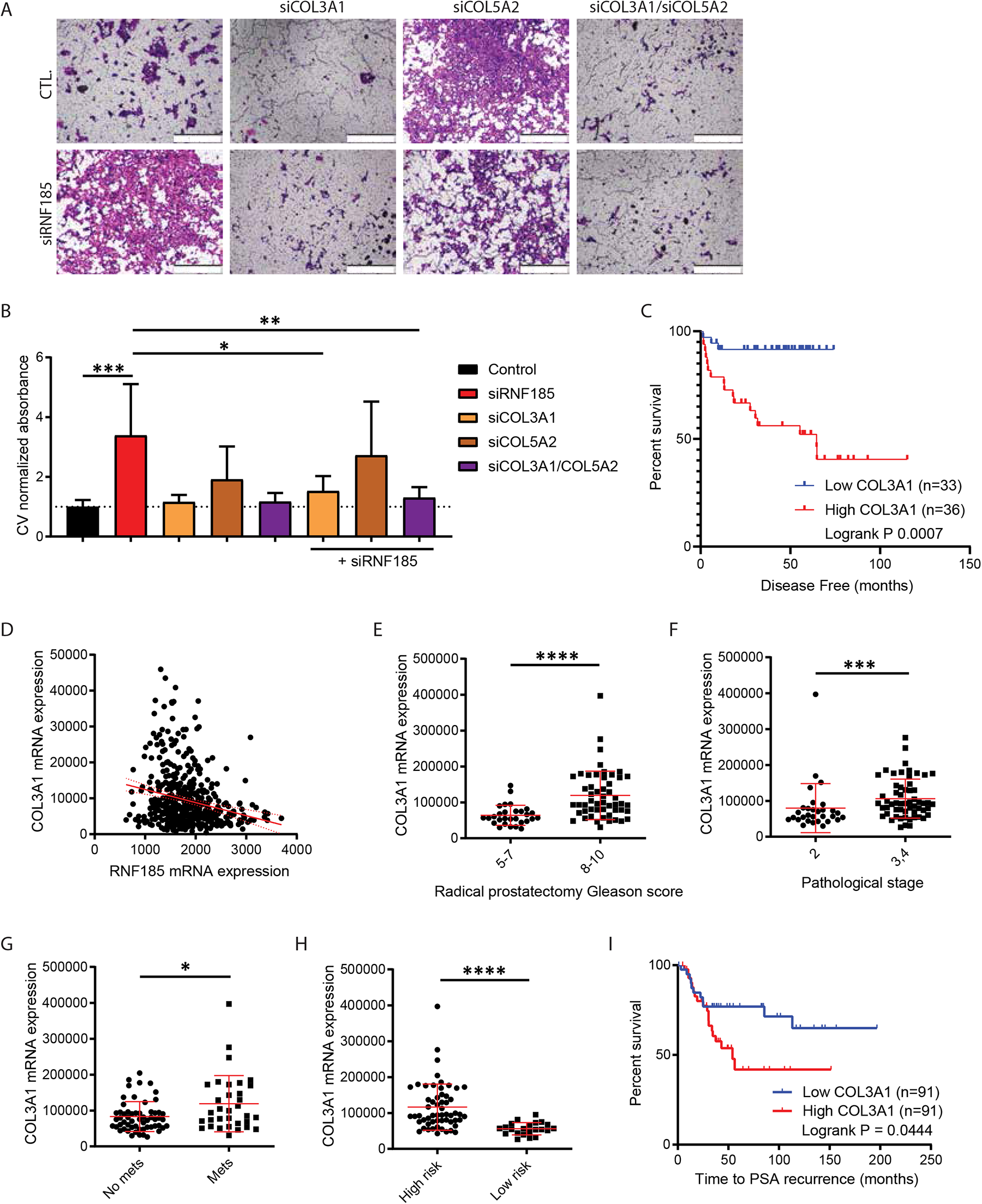
COL3A1 depletion in RNF185-deficient PC3 cells rescues migration advantage conferred by RNF185 KD alone. (A) Representative images of migrated PC3 cells transfected with the indicated siRNAs. (B) Quantification of migrated cells based on Crystal Violet (CV) absorbance. (C) Disease-free survival in samples stratified by COL3A1 mRNA expression (“Low COL3A1” = bottom quartile; “High COL3A1” = top quartile). (D) Scatter plot of RNF185 and COL3A1 mRNA expression in PCa patients (Pearson correlation=0.75, p-value=2.47e-53). (C) generated using datasets from Taylor et al., PMID: 20579941 (D) generated using TCGA PRAD dataset. Box plots show the median and whiskers (min to max), scatter plots show mean with SD. *P < 0.05, **P < 0.005, ***P < 0.001, ****P < 0.0001 by Mann-Whitney u test or one-way ANOVA. Survival graphs used the Kaplan–Meier with a log-rank (Mantel-Cox).

To validate these findings *in vivo*, we established stable MPC3 cells expressing shRNA against RNF185, COL3A1, and a combination of both (Supp. Fig. 5A-B). The increased migration and invasion phenotypes previously seen upon RFN185 KD, as well as the rescue of this phenotype in the RNF185/COL3A1 double KD, were confirmed in the Boyden chamber assay (Fig. 5A, Supp. Fig. 5C). Following this validation, RNF185 KD, COL3A1 KD, RNF185/COL3A1 double KD, and control MPC3 cells were inoculated subcutaneously in NSG mice. Analysis of tumor growth revealed significantly larger tumors in the RNF185 KD inoculated mice compared to the control group, as shown in earlier experiments (Fig. 2E, Supp. Fig.2H). Notably, COL3A1 KD in combination with RNF185 KD effectively attenuated the effect of RNF185 KD alone, as mice inoculated with RNF185/COL3A1 double KD cells grew tumors of comparable size as the control group or mice inoculated with the COL3A1 KD cells (Fig. 5B-F). Quantification of lung metastasis identified that a wider area of the lungs inoculated with RNF185 KD cells contained metastases, when compared to the mice inoculated with the control cells, consistent with our earlier findings (Fig. 2F). Notably, COL3A1 depletion was sufficient to reduce the metastatic load seen upon inoculation with RNF185 KD cells; mice inoculated with tumors that were inhibited for both RNF185 and COL3A1 exhibited a metastatic load which was comparable with that seen in control mice. Interestingly, COL3A1 depletion on its own did not affect the number of lung metastases (Fig. 5G). Taken together, these data suggest that COL3A1 upregulation underlies increased migration phenotype observed in RNF185-depleted cells, *in vitro* and *in vivo*, as depletion of COL3A1 is sufficient to attenuate the enhanced invasiveness of RNF185 KD prostate cancer cells.

**Fig. 5.**
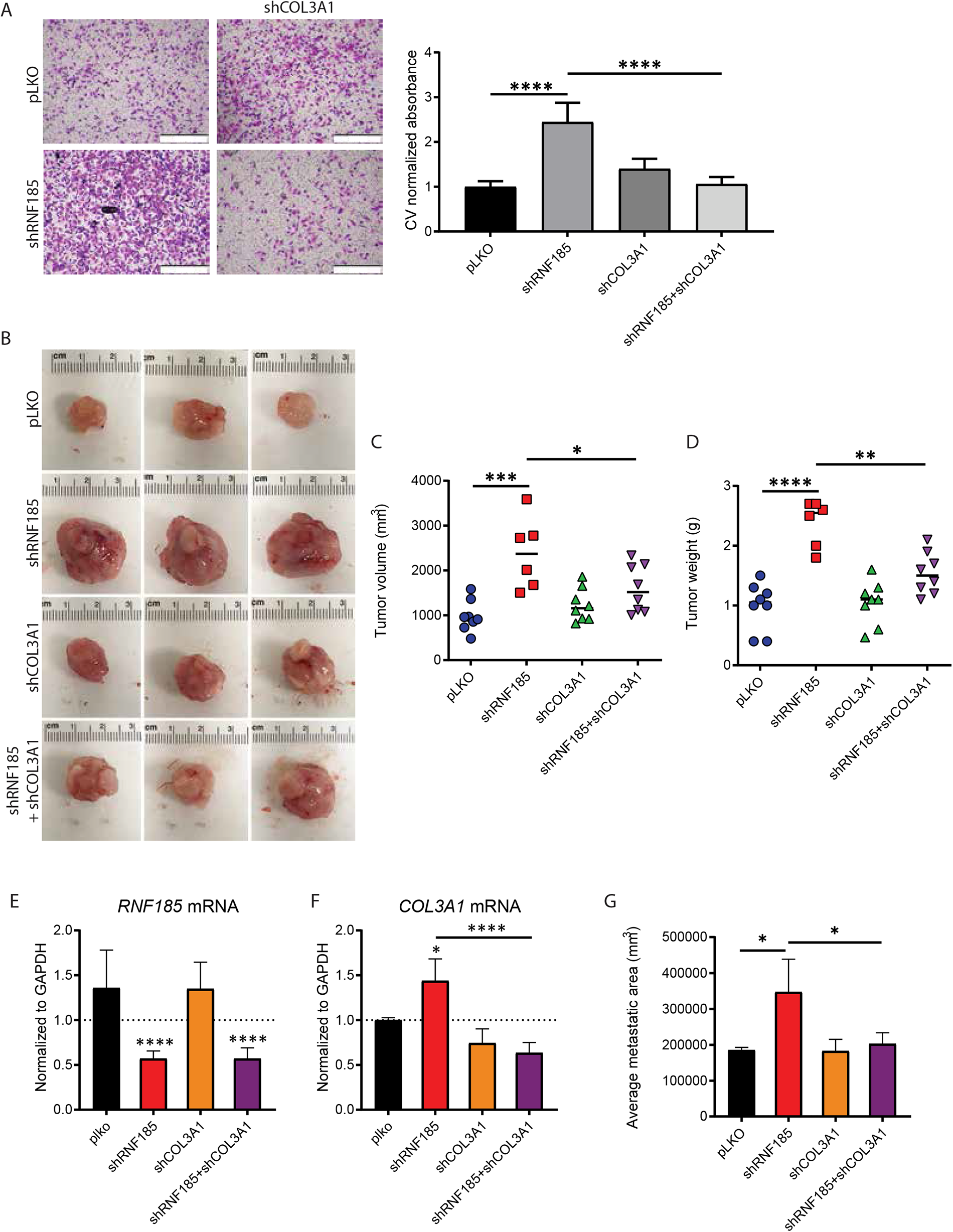
COL3A1 depletion in RNF185-deficient MPC3 cells rescues tumor growth advantage and metastatic propensity conferred by RNF185 KD alone in subcutaneous grafts. (A) Representative images of MPC3 cells expressing the indicated vectors that migrated through Matrigel and quantification of migrated cells based on Crystal Violet (CV) absorbance. (B) Representative pictures of resected tumors originating from cells expressing the indicated vectors. (C) Volume of resected tumors originating from cells expressing the indicated vectors 28 days after subcutaneous injection (n=6-8/group). (D) Weights of resected tumors originating from cells expressing the indicated vectors 28 days after subcutaneous injection (n=6-8/group). (E) qRT-PCR analysis of RNF185 mRNA in tumors emerging following subcutaneous injection of either MPC3.plko, MPC3.shRNF185, MPC3.COL3A1 or MPC3.shRNF185+shCOL3A1 cells into flanks of NSG mice. (F) qRT-PCR analysis of COL3A1 mRNA in tumors emerging following subcutaneous injection of either MPC3.plko, MPC3.shRNF185, MPC3.COL3A1 or MPC3.shRNF185+shCOL3A1 cells into flanks of NSG mice. (G) Average metastatic area (mm^3^) per section, measured in 9 serial sections of the lungs for each mouse within each group (n=6-8 mice per group). Scatter plots show the mean with SD, box plots show median with whiskers (min to max). *P < 0.05, **P < 0.005, ***P < 0.001, ****P < 0.0001 by one-way ANOVA.

## Discussion

AR-dependent signaling is commonly dysregulated in prostate cancer, which leads to transcriptional changes that promote cellular growth and to functional outcomes associated with neoplasia. These activities provide a rationale for targeting androgen signaling, a goal accomplished in part through ADT therapy. Unfortunately, ADT frequently rewires AR signaling pathways in a way that promotes therapy resistance. Prostate cancer that develops ADT resistance usually progresses to mCRPC, for which the effective treatment remains limited^52^. Here, we identify a role for the ubiquitin ligase RNF185 in prostate cancer progression and propensity to metastasize. Analyses of 3 independent prostate cancer patient datasets revealed that low RNF185 mRNA and protein expression was associated with poorer disease-free survival and higher Gleason grade and metastatic load. Notably, inhibition of RNF185 expression in mouse and human PCa lines increased migration and invasive phenotypes in culture and resulted in larger tumors and increased lung metastases *in vivo*. Correspondingly, gene expression analyses identified enrichment of genes associated with cell migration and the EMT and pointed to the role of specific collagens in phenotypes seen in RNF185 KD PCa cells. Notably, the changes elicited upon RNF185 KD were seen in some but not all PCa cell lines tested, suggesting that certain genetic alterations (as seen in mouse MPC3 or human PC3 and C4-2B prostate cancer cell lines) are prerequisites to confer phenotypic changes when combined with the loss of RNF185.

As a ubiquitin ligase, RNF185 was previously associated with cancer cell migration. Lower level of RNF185 expression was found in glioblastoma (GBM), which inversely correlated with prognosis^53^. RNF185 overexpression in a GBM cell line was sufficient to decrease cell migration *in vitro*^53^. On the other hand, RNF185 was shown to contribute to gastric cancer metastasis through ubiquitination and proteasomal degradation of JWA, a microtubule-associated protein involved in the DNA damage response^17^. Furthermore, RNF185 was identified as part of a ubiquitin-related gene signature that can be used to predict prostate cancer progression^54^. However, how RNF185 downregulation triggers changes that may result in enhanced metastasis was not known. Our data suggest that reduced RNF185 levels induce activities that enhance the EMT, which in turn impacts metastasis. DEG analysis in RNF185-depleted, versus control, conditions following transcriptional mapping of clinical samples and RNAseq analysis of mouse prostate cancer lines identified genes implicated in cell adhesion, cytoskeleton remodeling, ECM structural regulation, and cancer cell migration. Among those, COL5A2 and COL3A1 are reportedly often overexpressed in cancers^21, 22^. Collagen is a key component of the tumor microenvironment; thus, collagen dysregulation may underlie the tumor-intrinsic propensity to metastasize. COL5A2 was previously shown to promote proliferation and migration in prostate cancer lines, and its levels positively correlate with Gleason grade in PCa patients^55^. COL5A2 is also upregulated and involved in tumor progression in colorectal cancer^56^. COL3A1 was found to be a prognostic biomarker for ovarian carcinoma, breast cancer, and colorectal carcinoma^25, 26, 55^. Our studies reveal that upregulation of COL3A1 alone is sufficient to drive enhanced migration phenotypes seen in mouse and human prostate cancer cells.

How RNF185 controls COL3A1 expression remains to be determined. Our observations suggest a cross talk between COL3A1 and COL5A2 transcription. Both genes are located within the same gene cluster on chromosome 2 and are often co-expressed, as well as deregulated in connective tissue diseases^58–60^. Their transcription has been reported to be controlled by TGF-beta signaling, which has been implicated in metastasis of advanced stages of prostate cancer^61–64^. This implies that TGF-beta may be involved in the regulatory changes induced by RNF185, a possibility that warrant further exploration. Correspondingly, upregulated COL3A1 expression coincides with decreased RNF185 expression in metastatic PCa, substantiating the clinical relevance of this newly identified regulatory axis for PCa progression and metastasis. Our results show that manipulating COL3A1 and RNF185 expression is sufficient to impact prostate tumor cells migration and suggest that components of this regulatory pathway may serve as markers to predict PCa outcome. Further, as inhibition of COL3A1 attenuates metastasis in cells with reduced RNF185 expression, targeting collagen becomes an appealing consideration, which may provide the basis for future therapeutic evaluations in low RNF185 expressing advanced prostate tumors.

## Author Contributions

BVE and ZAR designed the study. BVE performed the experiments. AM performed pathological assessments. LF, HZO and RM performed bioinformatic analyses. MG provided clinical samples. BVE and ZAR wrote the manuscript.

## Acknowledgements

We thank members of the Ronai lab for discussion. We thank SBP shared resources, which is supported by P30CA030199 grant, for their help with the analyses (vivarium, FACS, and bioinformatics). Support by R35CA197465 (to ZAR) is gratefully acknowledged.

## Supplementary Figure Legends

**Supplementary Fig. 1.**
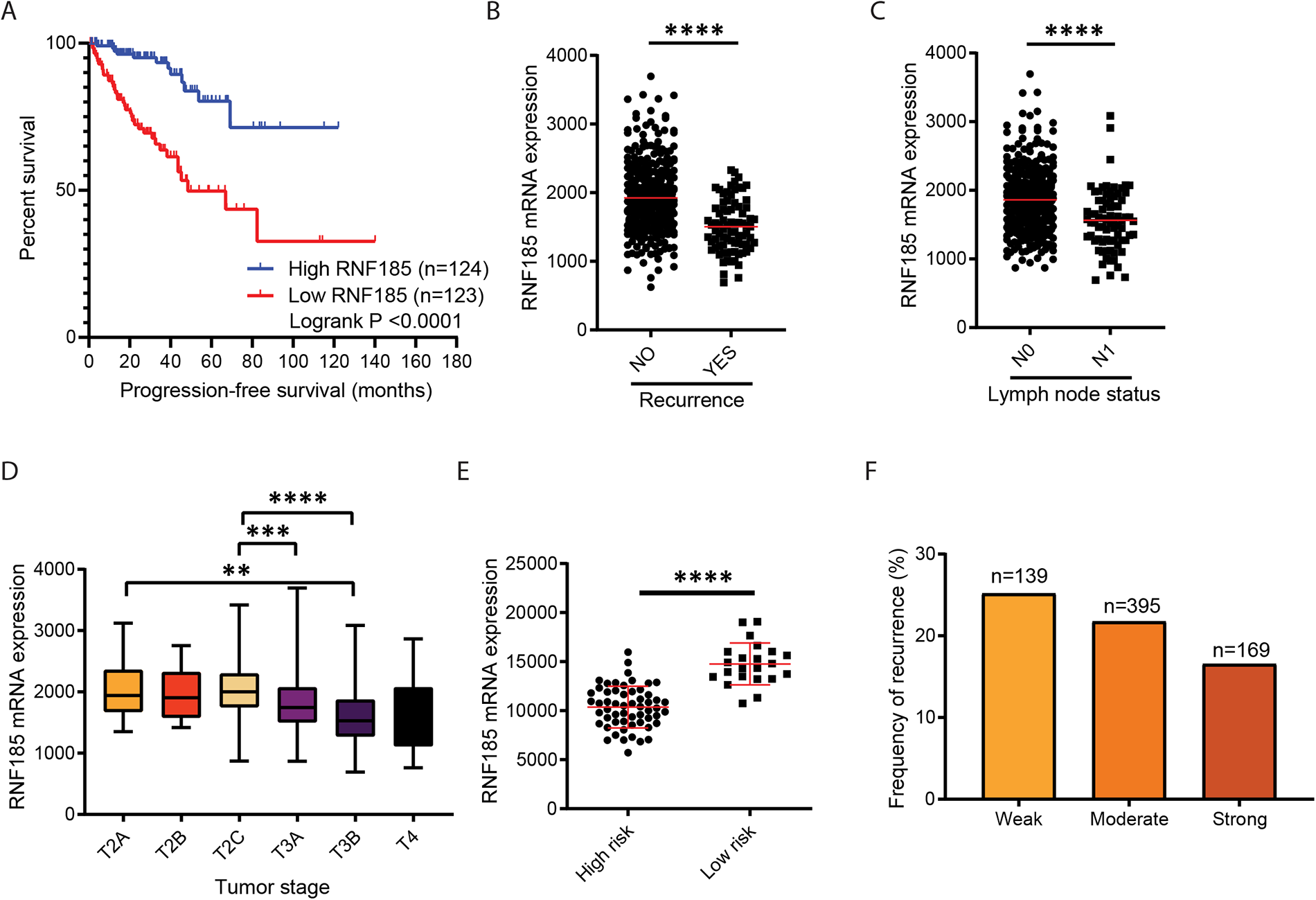
RNF185 expression correlates with PCa patient outcome. (A) Progression-free survival in samples stratified based on RNF185 mRNA expression (“Low RNF185” = bottom quartile; “High RNF185” = top quartile). (B) RNA-seq analysis of RNF185 in samples sorted by New Neoplasm Event post initial therapy (NO: n=351; YES: n=87). (C) RNA-seq analysis of RNF185 in samples sorted by New Neoplasm Event post initial therapy (N0: n=338; N1: n=77). (D) RNA-seq analysis of RNF185 mRNA in samples sorted by Tumor stage (T2A: n=13; T2B: n=10; T2C: n=163; T3A: n=157; T3B: n=133; T4: n=10). (E) RNA-seq analysis of RNF185 in samples sorted by risk (High: n=57; Low: n=22) (F) Recurrence percentage in PCa TMAs stratified according to RNF185 IHC staining intensity. (A-D) generated using TCGA’s Pan-Cancer Atlas and Firehose legacy PRAD datasets. (E-F) generated from VPC unpublished cohorts. Box plots and scatter plots show the median and whiskers (min to max). *P < 0.05, **P < 0.005, ***P < 0.001, ****P < 0.0001 by two-tailed t-test or one-way ANOVA. Survival graphs used the Kaplan–Meier with a log-rank (Mantel-Cox).

**Supplementary Fig. 2.**
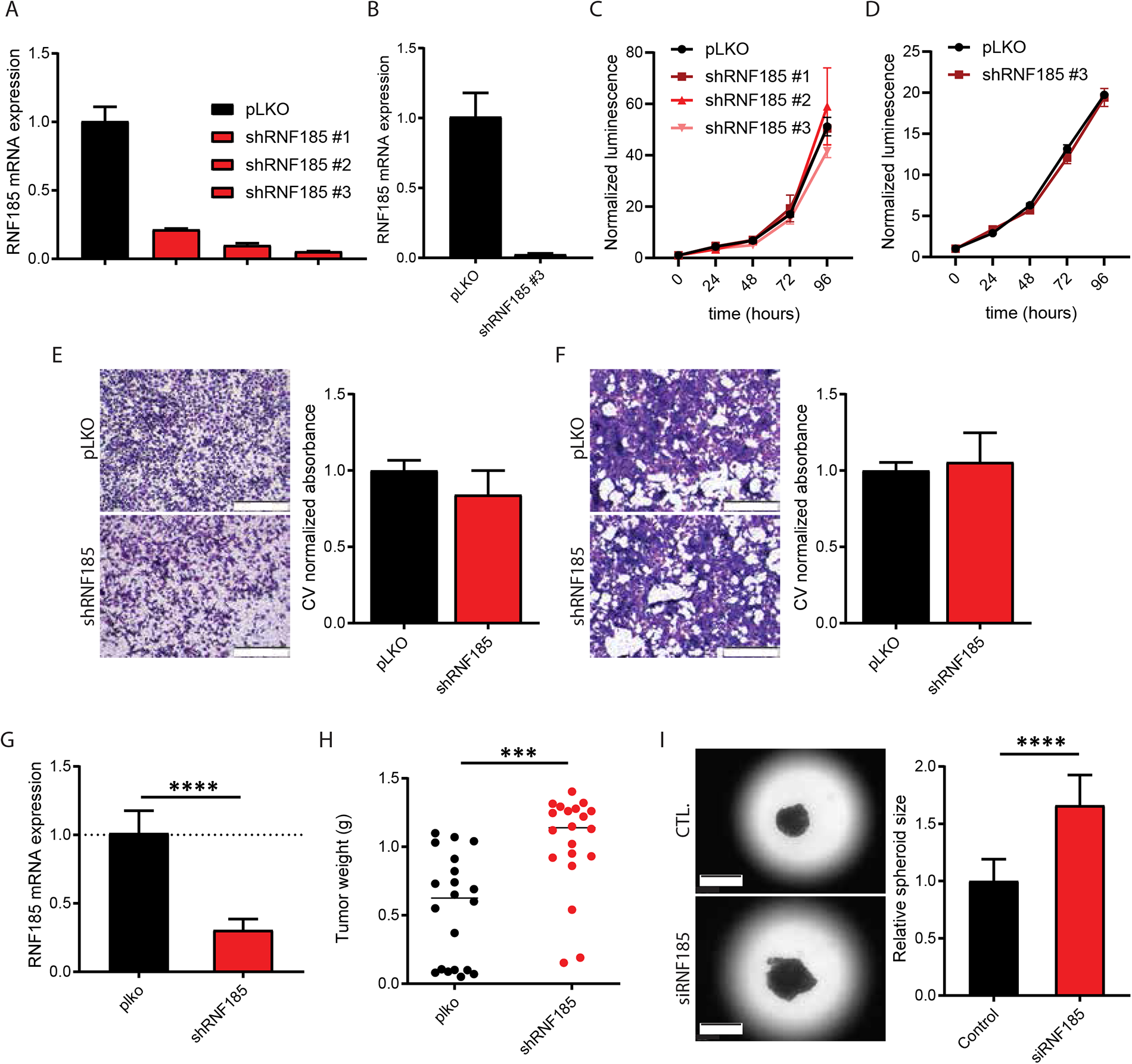
Effect of RNF185 KD in PCa cell lines and validation of *in vivo* phenotype. (A) qRT-PCR analysis of RNF185 mRNA in MyC-CaP cells expressing the indicated vectors. (B) qRT-PCR analysis of RNF185 mRNA in TRAMP-C2 cells expressing the indicated vectors. Representative pictures of migrated MPC3 cells expressing the indicated vectors and quantification of those migrated cells based on Crystal Violet (CV) absorbance. (C) Proliferation assay of cells shown in (A). (D) Proliferation assay of cells shown in (B). (E) Representative images of migrated MyC-CaP cells expressing the indicated vectors and quantification of migrated cells based on Crystal Violet (CV) absorbance. (F) Representative images of migrated TRAMP-C2 cells expressing the indicated vectors and quantification of migrated cells based on Crystal Violet (CV) absorbance. (G) qRT-PCR analysis of RNF185 mRNA in tumors emerging following subcutaneous injection of either MPC3.plko or MPC3.shRNF185 cells into flanks of NSG mice. (H) Weight of resected tumors originating from MPC3-pLKO or MPC3-shRNF185 cells 34 days after subcutaneous injection (n=20/group). (I) Representative pictures of spheroids from C4-2B cells transfected with the indicated siRNA after 9 days in culture. (J) Quantification of relative spheroid size compared to controls (n=24 spheroids/group) measured by surface area with ImageJ. Box plots show the median and whiskers (min to max). Growth curve shows the mean and whiskers (min to max). Scatter plot shows all values and median. ***P < 0.001, ****P < 0.0001 by two-tailed t-test.

**Supplemental Fig. 3.**
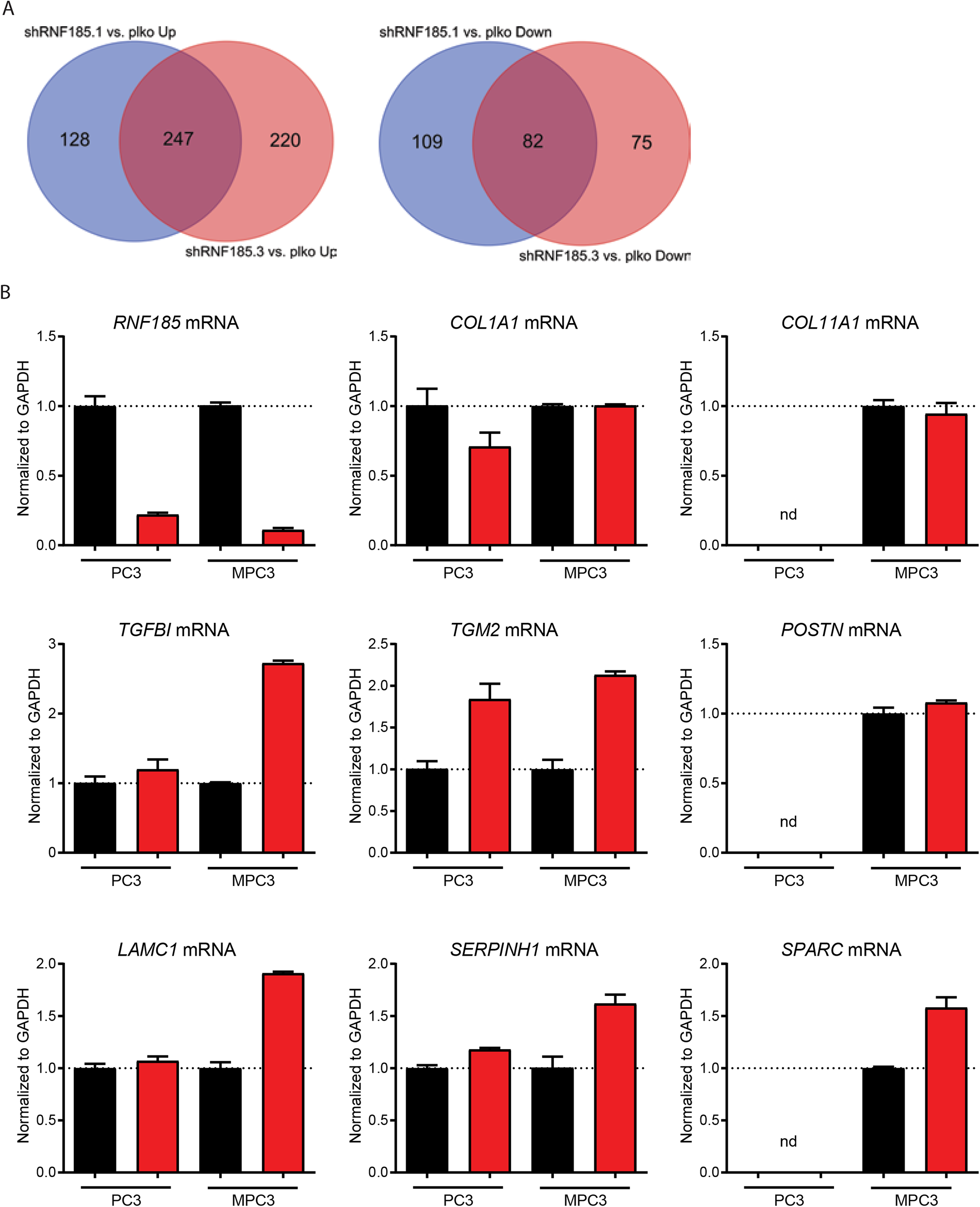
RT-qPCR validation of genes that are differentially expressed in both RNF185 KD MPC3 and PC3 cells. (A) Venn Diagrams of upregulated and downregulated genes in MPC3-shRNF185 #1 and #3 cells. (B) qRT-PCR analysis of indicated genes in MPC3-pLKO and MPC3-shRNF185 cells, and in the human prostate cancer line PC3 transfected with RNF185 siRNA. Box plots show median and whiskers (nd = not detected).

**Supplementary Fig. 4.**
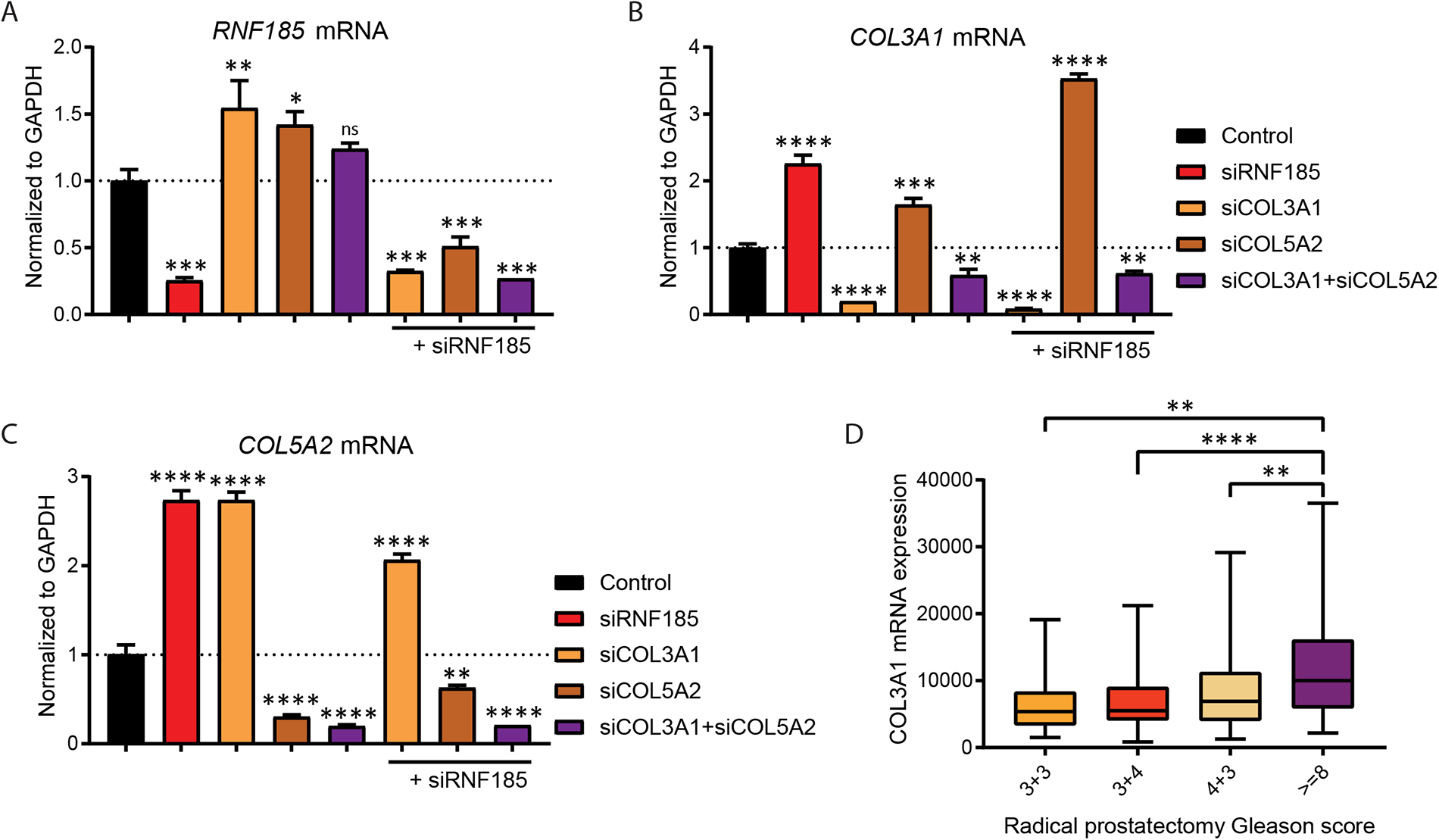
RT-qPCR validation of RNF185, COL3A1 and COL5A2 expression in RNF185 and COL3A1 KD PC3 cells. qRT-PCR analysis of RNF185 (A), COL3A1 (B) and COL5A2 transcripts in PC3 cells transfected with indicated siRNAs. (D) RNA-seq analysis of RNF185 mRNA in samples sorted by Gleason score (3+3: n=19; 3+4: n=74; 4+3: n=43; >=8: n=98). (D) generated using datasets from Taylor et al., PMID: 20579941. Box plots show median and whiskers. *P < 0.05, **P < 0.005, ***P < 0.001, ****P < 0.0001 by one-way ANOVA. Survival graphs used the Kaplan–Meier with a log-rank (Mantel-Cox).

**Supplementary Fig. 5.**
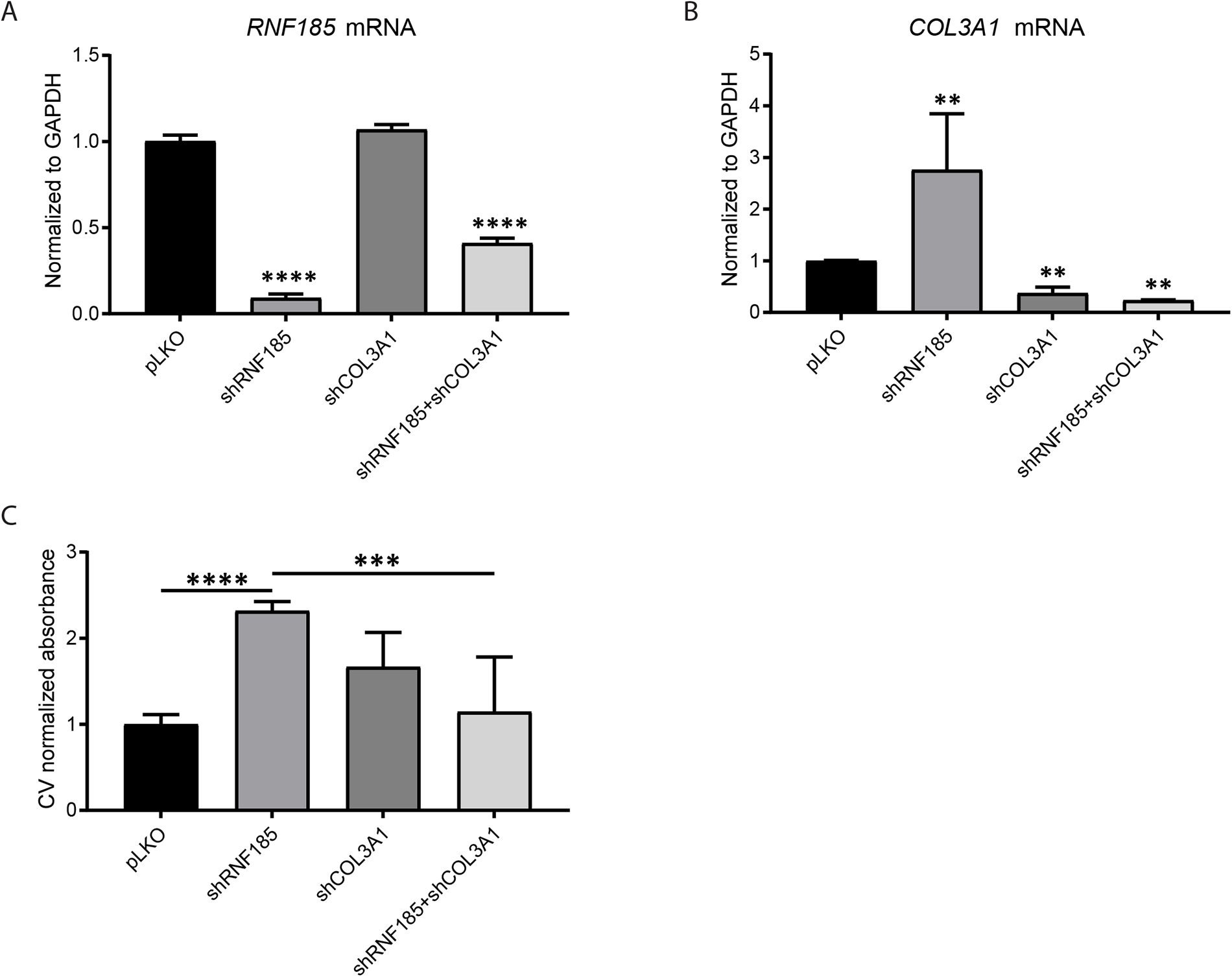
RT-qPCR validation of RNF185 and COL3A1 in stable RNF185 and COL3A1 KD MPC3 cells. qRT-PCR analysis of RNF185 (A), COL3A1 (B) transcripts in MPC3 cells expressing the indicated shRNAs. (C) Quantification of migrated MPC3 cells expressing the indicated vectors based on Crystal Violet (CV) absorbance in Boyden chamber assay. Box plots show median and whiskers. **P < 0.005, ***P < 0.001, ****P < 0.0001 by one-way ANOVA.

## Supplemental Tables

**Supplemental Table 1:** GSEA analysis identifies genes that are enriched in PRAD patients (HALLMARL_EMT in TCGA PRAD) with low RNF185 expression (bottom quartile) compared to patients with high RNF185 expression (top quartile).

**Supplemental Table 2:** GSEA analysis identifies genes that are enriched upon inhibition of RNF185 expression in MPC3 cells (HALLMARL_EMT in MPC3.shRNF185#1 compared to MPC3.pLKO)

**Supplemental Table 3:** GSEA analysis identifies genes that are enriched upon inhibition of RNF185 expression in MPC3 cells (HALLMARL_EMT in MPC3.shRNF185#3 compared to MPC3.pLKO)

