## supplemental tables 1-3 for "RNF185 control of COL3A1 expression limits prostate cancer migration and metastatic potential"

**Supplemental Table 1:** **GSEA analysis identifies genes that are enriched in PRAD patients (HALLMARL_EMT in TCGA PRAD) with low RNF185 expression (bottom quartile) compared to patients with high RNF185 expression (top quartile).**

| SYMBOL | RANK IN GENE LIST | RANK METRIC SCORE | RUNNING ES | CORE ENRICHMENT |
| --- | --- | --- | --- | --- |
| CTHRC1 | 3 | 40.16309 | 0.057418 | Yes |
| COMP | 4 | 39.72491 | 0.114361 | Yes |
| COL11A1 | 30 | 30.1932 | 0.156357 | Yes |
| FAP | 44 | 27.099 | 0.194534 | Yes |
| SFRP4 | 53 | 24.3443 | 0.229019 | Yes |
| THBS2 | 95 | 19.30602 | 0.254587 | Yes |
| BGN | 128 | 16.33554 | 0.276359 | Yes |
| INHBA | 159 | 14.90255 | 0.29618 | Yes |
| COL1A1 | 161 | 14.80937 | 0.317357 | Yes |
| THY1 | 201 | 12.29829 | 0.332982 | Yes |
| MATN3 | 284 | 9.656747 | 0.342612 | Yes |
| VCAN | 300 | 9.280169 | 0.355144 | Yes |
| CRLF1 | 374 | 8.11612 | 0.363027 | Yes |
| COL5A2 | 379 | 8.06271 | 0.374379 | Yes |
| NTM | 384 | 8.015413 | 0.385663 | Yes |
| SPP1 | 400 | 7.775103 | 0.396038 | Yes |
| MCM7 | 474 | 6.867 | 0.402131 | Yes |
| ADAM12 | 508 | 6.457201 | 0.409691 | Yes |
| GREM1 | 557 | 6.080195 | 0.415941 | Yes |
| IGFBP3 | 579 | 5.910374 | 0.423334 | Yes |
| MMP3 | 673 | 5.195263 | 0.426003 | Yes |
| SERPINE1 | 690 | 5.075 | 0.432456 | Yes |
| MGP | 735 | 4.869651 | 0.437176 | Yes |
| COL3A1 | 777 | 4.672551 | 0.441767 | Yes |
| SERPINH1 | 886 | 4.155785 | 0.442176 | Yes |
| SPARC | 917 | 4.055232 | 0.446447 | Yes |
| LOX | 964 | 3.907054 | 0.449685 | Yes |
| POSTN | 987 | 3.838395 | 0.454056 | Yes |
| IL32 | 1231 | 3.164565 | 0.446108 | Yes |
| TNFRSF12A | 1251 | 3.125206 | 0.449612 | Yes |
| MEST | 1342 | 2.916188 | 0.449168 | Yes |
| TGM2 | 1366 | 2.872128 | 0.452103 | Yes |
| LAMC1 | 1369 | 2.870705 | 0.456116 | Yes |
| TGFB1 | 1392 | 2.808231 | 0.459011 | Yes |
| NNMT | 1515 | 2.575925 | 0.456435 | Yes |
| PLAUR | 1534 | 2.54042 | 0.459152 | Yes |
| COL1A2 | 1551 | 2.520354 | 0.461943 | Yes |
| ELN | 1684 | 2.308 | 0.458469 | Yes |
| TGFBI | 1720 | 2.258921 | 0.459909 | Yes |
| SERPINE2 | 1743 | 2.211875 | 0.46195 | Yes |
| FZD8 | 1755 | 2.190632 | 0.464525 | Yes |

**Supplemental Table 2: GSEA analysis identifies genes that are enriched upon inhibition of RNF185 expression in MPC3 cells (HALLMARL_EMT in MPC3.shRNF185#1 compared to MPC3.pLKO).**

| SYMBOL | RANK IN GENE LIST | RANK METRIC SCORE | RUNNING ES | CORE ENRICHMENT |
| --- | --- | --- | --- | --- |
| Col11a1 | 19 | 2.798753 | 0.023755 | Yes |
| Fbln5 | 21 | 2.716667 | 0.047961 | Yes |
| Loxl1 | 55 | 2.206156 | 0.0655 | Yes |
| Myl9 | 59 | 2.183487 | 0.084811 | Yes |
| Serpinh1 | 86 | 2.047921 | 0.101397 | Yes |
| Postn | 102 | 1.970229 | 0.118013 | Yes |
| Tgfbi | 219 | 1.530797 | 0.124058 | Yes |
| Sntb1 | 235 | 1.498852 | 0.136462 | Yes |
| Snai2 | 250 | 1.482931 | 0.14879 | Yes |
| Slit2 | 251 | 1.482499 | 0.162035 | Yes |
| Ptx3 | 263 | 1.46303 | 0.174383 | Yes |
| Col3a1 | 289 | 1.404287 | 0.185284 | Yes |
| Col5a3 | 296 | 1.394408 | 0.197348 | Yes |
| Col12a1 | 303 | 1.389293 | 0.209365 | Yes |
| Gas1 | 340 | 1.337846 | 0.21895 | Yes |
| Tgm2 | 367 | 1.294192 | 0.228802 | Yes |
| Col5a1 | 380 | 1.269931 | 0.239358 | Yes |
| Lamc1 | 395 | 1.254988 | 0.24965 | Yes |
| Timp3 | 397 | 1.251897 | 0.260769 | Yes |
| Col16a1 | 428 | 1.221998 | 0.269713 | Yes |
| Serpine1 | 480 | 1.161287 | 0.276733 | Yes |
| Mmp2 | 486 | 1.155214 | 0.286725 | Yes |
| Msx1 | 498 | 1.139903 | 0.296185 | Yes |
| Col6a2 | 508 | 1.125452 | 0.305648 | Yes |
| Matn2 | 511 | 1.121436 | 0.315536 | Yes |
| Fas | 527 | 1.105198 | 0.324423 | Yes |
| Tgfbr3 | 529 | 1.101545 | 0.334199 | Yes |
| Fmod | 531 | 1.099167 | 0.343954 | Yes |
| Fstl1 | 539 | 1.09076 | 0.353238 | Yes |
| Qsox1 | 543 | 1.086996 | 0.362753 | Yes |
| Pmp22 | 557 | 1.071967 | 0.371475 | Yes |
| Cxcl5 | 611 | 1.020204 | 0.377102 | Yes |
| Col1a1 | 612 | 1.019844 | 0.386214 | Yes |
| Spock1 | 696 | 0.951497 | 0.389254 | Yes |
| Ecm1 | 701 | 0.95051 | 0.397483 | Yes |
| Fbln1 | 764 | 0.906308 | 0.401501 | Yes |
| Tnc | 774 | 0.901049 | 0.408959 | Yes |
| Prrx1 | 797 | 0.887692 | 0.415443 | Yes |
| Col5a2 | 805 | 0.881753 | 0.42286 | Yes |
| Sgcb | 815 | 0.876359 | 0.430098 | Yes |
| Dst | 837 | 0.865741 | 0.436451 | Yes |
| Eno2 | 966 | 0.794017 | 0.435124 | Yes |
| Thbs2 | 970 | 0.792622 | 0.442008 | Yes |
| Il6 | 1073 | 0.750048 | 0.441998 | Yes |
| Sdc1 | 1085 | 0.744256 | 0.447924 | Yes |
| Sparc | 1127 | 0.72144 | 0.451672 | Yes |
| Lox | 1194 | 0.69936 | 0.453578 | Yes |
| Cxcl12 | 1208 | 0.694605 | 0.458928 | Yes |
| Plod1 | 1249 | 0.678696 | 0.46236 | Yes |
| Wipf1 | 1297 | 0.661085 | 0.465174 | Yes |
| Itga5 | 1309 | 0.656968 | 0.47032 | Yes |
| Lrp1 | 1428 | 0.623871 | 0.46813 | Yes |
| Pdgfrb | 1445 | 0.618318 | 0.472602 | Yes |
| P3h1 | 1495 | 0.605767 | 0.47479 | Yes |
| Itgb5 | 1638 | 0.566397 | 0.470508 | Yes |
| Lama2 | 1668 | 0.559652 | 0.4736 | Yes |
| Lrrc15 | 1709 | 0.551553 | 0.475896 | Yes |
| Acta2 | 1714 | 0.549957 | 0.480546 | Yes |
| Sgcd | 1824 | 0.522426 | 0.478042 | Yes |
| Pcolce | 1923 | 0.502099 | 0.47608 | Yes |
| Jun | 2022 | 0.483551 | 0.473953 | Yes |
| Plod3 | 2108 | 0.467454 | 0.472537 | Yes |
| Gem | 2133 | 0.463019 | 0.475095 | Yes |
| Col4a1 | 2173 | 0.455039 | 0.476594 | Yes |
| Fbln2 | 2217 | 0.446697 | 0.477756 | Yes |
| Col6a3 | 2265 | 0.439828 | 0.478593 | Yes |
| Magee1 | 2287 | 0.436635 | 0.481113 | Yes |
| Grem1 | 2322 | 0.430092 | 0.482718 | Yes |

**Supplemental Table 3: GSEA analysis identifies genes that are enriched upon inhibition of RNF185 expression in MPC3 cells (HALLMARL_EMT in MPC3.shRNF185#3 compared to MPC3.pLKO)**

| SYMBOL | RANK IN GENE LIST | RANK METRIC SCORE | RUNNING ES | CORE ENRICHMENT |
| --- | --- | --- | --- | --- |
| Sntb1 | 42 | 2.469361 | 0.01961 | Yes |
| Fmod | 46 | 2.439663 | 0.041516 | Yes |
| Col3a1 | 50 | 2.398955 | 0.063054 | Yes |
| Tgfbi | 75 | 2.221706 | 0.081604 | Yes |
| Lrrc15 | 83 | 2.167979 | 0.100785 | Yes |
| Snai2 | 114 | 2.008835 | 0.117012 | Yes |
| Col1a1 | 159 | 1.879458 | 0.131145 | Yes |
| Col12a1 | 186 | 1.803005 | 0.14577 | Yes |
| Myl9 | 206 | 1.741683 | 0.1603 | Yes |
| Col6a2 | 215 | 1.707047 | 0.17524 | Yes |
| Col5a2 | 240 | 1.652353 | 0.188632 | Yes |
| Col6a3 | 313 | 1.511696 | 0.197591 | Yes |
| Gas1 | 328 | 1.495297 | 0.210217 | Yes |
| Col11a1 | 383 | 1.419756 | 0.219528 | Yes |
| Col5a1 | 396 | 1.388735 | 0.23132 | Yes |
| Fbln5 | 475 | 1.270301 | 0.237698 | Yes |
| Cxcl12 | 482 | 1.264408 | 0.248759 | Yes |
| Ecm1 | 487 | 1.259289 | 0.259905 | Yes |
| Mylk | 551 | 1.176078 | 0.266415 | Yes |
| Sparc | 570 | 1.164008 | 0.275777 | Yes |
| Tgm2 | 585 | 1.156454 | 0.285334 | Yes |
| Col16a1 | 587 | 1.15508 | 0.295733 | Yes |
| Fstl1 | 605 | 1.142236 | 0.304963 | Yes |
| Acta2 | 621 | 1.122094 | 0.314143 | Yes |
| Itga5 | 627 | 1.119772 | 0.323959 | Yes |
| Matn2 | 628 | 1.119478 | 0.334102 | Yes |
| Eno2 | 633 | 1.112347 | 0.343917 | Yes |
| Loxl1 | 678 | 1.081064 | 0.350817 | Yes |
| Lama2 | 679 | 1.080235 | 0.360604 | Yes |
| Cxcl5 | 739 | 1.039234 | 0.366138 | Yes |
| Pmp22 | 790 | 0.999534 | 0.371904 | Yes |
| Pdgfrb | 796 | 0.99517 | 0.380591 | Yes |
| Postn | 835 | 0.96844 | 0.386865 | Yes |
| Edil3 | 838 | 0.967032 | 0.395495 | Yes |
| Serpinh1 | 840 | 0.964879 | 0.404172 | Yes |
| Prrx1 | 873 | 0.945646 | 0.410634 | Yes |
| Il15 | 877 | 0.943921 | 0.418989 | Yes |
| Col1a2 | 893 | 0.931563 | 0.426442 | Yes |
| Tgfbr3 | 897 | 0.928283 | 0.434655 | Yes |
| Tnfrsf11b | 906 | 0.925235 | 0.442511 | Yes |
| Lum | 925 | 0.913446 | 0.449603 | Yes |
| Fbn2 | 948 | 0.8981 | 0.456292 | Yes |
| Lamc1 | 964 | 0.888828 | 0.463359 | Yes |
| Ptx3 | 970 | 0.887054 | 0.471066 | Yes |
| Tpm2 | 1059 | 0.831924 | 0.472814 | Yes |
| Bgn | 1065 | 0.827029 | 0.479978 | Yes |
| Tnc | 1138 | 0.795238 | 0.482446 | Yes |
| Tagln | 1194 | 0.773542 | 0.485836 | Yes |
| Rgs4 | 1251 | 0.7505 | 0.488951 | Yes |
| Lrp1 | 1287 | 0.741002 | 0.493362 | Yes |
| P3h1 | 1304 | 0.734463 | 0.498963 | Yes |
| Qsox1 | 1324 | 0.730009 | 0.504327 | Yes |
| Plod1 | 1356 | 0.718045 | 0.508793 | Yes |
| Cdh6 | 1407 | 0.703529 | 0.511878 | Yes |
| Fn1 | 1419 | 0.699849 | 0.517495 | Yes |
| Sdc1 | 1464 | 0.683248 | 0.52079 | Yes |
| Lama3 | 1541 | 0.660904 | 0.521778 | Yes |
| Sgcb | 1605 | 0.639595 | 0.523428 | Yes |
| Pdlim4 | 1759 | 0.595028 | 0.518752 | Yes |
| Timp3 | 1761 | 0.594673 | 0.524074 | Yes |
| Mmp2 | 2020 | 0.531908 | 0.511919 | Yes |
| Mmp14 | 2062 | 0.520509 | 0.513937 | Yes |
| Fermt2 | 2078 | 0.516244 | 0.517628 | Yes |
| Plod2 | 2161 | 0.498963 | 0.516753 | Yes |
| Cdh11 | 2259 | 0.479504 | 0.514715 | Yes |
| Fbln1 | 2281 | 0.476458 | 0.517651 | Yes |
| Itgav | 2321 | 0.46701 | 0.519316 | Yes |
| Glipr1 | 2374 | 0.457304 | 0.520038 | Yes |
| Jun | 2438 | 0.448018 | 0.519952 | Yes |
| Efemp2 | 2453 | 0.445845 | 0.52307 | Yes |
| Serpine2 | 2498 | 0.439202 | 0.524155 | Yes |
| Fstl3 | 2505 | 0.438628 | 0.527734 | Yes |
| Col5a3 | 2635 | 0.415164 | 0.523008 | Yes |
| Lox | 2712 | 0.403474 | 0.521663 | Yes |
| Thbs2 | 2713 | 0.403345 | 0.525318 | Yes |
| Plod3 | 2767 | 0.394606 | 0.525406 | Yes |
| Anpep | 2768 | 0.39445 | 0.52898 | Yes |
| Il6 | 2785 | 0.392156 | 0.53148 | Yes |
| Grem1 | 2787 | 0.391974 | 0.534965 | Yes |
| Gpc1 | 2896 | 0.374966 | 0.531257 | Yes |
| Fas | 2910 | 0.371807 | 0.53377 | Yes |
| Pcolce | 3062 | 0.349058 | 0.526998 | Yes |
| Col4a2 | 3089 | 0.344658 | 0.52841 | Yes |
| Magee1 | 3135 | 0.338853 | 0.528519 | Yes |
| Dst | 3139 | 0.337694 | 0.531381 | Yes |
| Col7a1 | 3147 | 0.336737 | 0.533972 | Yes |
| Flna | 3170 | 0.333764 | 0.535548 | Yes |
